# Entomological investigations into yellow fever outbreak in northern Ghana

**DOI:** 10.1101/2024.11.18.624057

**Authors:** Joseph Harold Nyarko Osei, Sellase Pi-Bansa, Kwadwo Kyeremeh Frempong, Mavis Ofei, Helena Anokyewaa Boakye, Jane Ansah-Owusu, Sandra-Candys Adwirba Akorful, Richard Odoi-Teye Malm, Christopher Nii Laryea Tawiah-Mensah, Mufeez Abudu, Andy Asafu-Adjaye, Seth Offei Addo, Bright Agbodzi, Ronald Bentil, Deborah Pratt, Shirley Nimo-Paintsil, Joseph Humphrey Kofi Bonney, Maxwell Alexander Appawu, Millie-Cindy Aba Aude Koffi, Sylvester Coleman, Millicent Captain-Esoah, Chrysantus Kubio, Daniel Adjei Boakye, Samuel Kweku Dadzie

**Affiliations:** Department of Parasitology, Noguchi Memorial Institute for Medical Research, College of Health Sciences, University of Ghana, Legon, Ghana; Department of Virology, Noguchi Memorial Institute for Medical Research, College of Health Sciences, University of Ghana, Legon, Ghana; Department of Animal Biology and Conservation Science, College of Basic and Applied Sciences, University of Ghana, Legon, Ghana; Department of Biological, Environmental and Occupational Health, School of Public Health, College of Health Sciences, University of Ghana, Legon, Ghana; Acute Febrile Laboratory, Naval Medical Research Unit 3, Noguchi Memorial Institute for Medical Research, College of Health Sciences, University of Ghana, Legon, Ghana; Department of Theoretical and Applied Biology, College of Science, Kwame Nkrumah University of Science and Technology, Kumasi, Ghana; Department of Molecular Medicine, School of Medicine and Dentistry, Kwame Nkrumah University of Science and Technology, Kumasi, Ghana; Biomedical and Public Health Research Unit, Water Research Institute, Council for Scientific and Industrial Research, Accra, Ghana; Liverpool School of Tropical Medicine, Liverpool, United Kingdom; Department of Applied Biology, C. K. Tedam University of Technology and Applied Sciences, Navrongo, Ghana; Savannah Regional Health Directorate, Damongo, Ghana

**Keywords:** Aedes aegypti, Chikungunya, Dengue, Yellow Fever, Zika, outbreak, epidemic

## Abstract

**Background:** Recently, arboviruses have been of concern as pathogens for emerging and re-emerging infectious diseases. In Ghana, yellow fever outbreak occurred in Savannah Region in the year 2021. A team from different institutions, organisations, and stakeholders of health with varying vital expertise was assembled to respond to this national emergency to assess, contain and/or control the rapid spread of the disease. This paper presents findings from the entomological investigations conducted during the yellow fever outbreak.

**Methods:** Immature stages of *Aedes* mosquitoes were collected from breeding containers in and around houses, and adult mosquitoes sampled using BG-Sentinel traps, human landing catches and Prokopack collections. After morphological identification of these mosquitoes, they were screened for Chikungunya, Dengue Fever, Yellow Fever, and Zika viruses using real time reverse-transcription polymerase chain reaction.

**Results:** In all, 12,264 breeding containers were examined. A total of 3,885 larvae and 1,186 pupae were obtained from 173 containers. Out of 1,001 houses surveyed, 130 were positive for larvae and/or pupae. The breeding receptacles included plastic (6,529), metallic (6,024), clay jar (753), tire (565), and well (34). The WHO thresholds for arboviruses larval indices were used to assess risk. A total of 1571 adults identified [*Aedes aegypti aegypti* (35), *Aedes aegypti formosus* (619), and *Culex* (917)] were collected with adult mosquito sampling methods or emerged from immature mosquitoes stages. None of the arboviruses were detected using qPCR.

**Conclusion:** Vectors had no yellow fever infections. There was a high risk of arbovirus transmission in the study areas although mosquito vectors were not positive for arboviruses. *Aedes aegypti formosus* was the dominant *Aedes* species. They might be drivers for yellow fever transmission during outbreak. Generally, arboviral transmission was high in all study districts. Although yellow fever virus was not detected, *Aedes aegypti* populations and transmission risk in study districts was high.

**Author summary:** In 2021, Savannah Region of Ghana experienced yellow fever outbreak. This spread quickly to adjoining regions. The disease is transmitted by some female mosquitoes infected with yellow fever virus. These same mosquitoes can transmit other viral infections resulting in disease outcomes such as chikungunya, dengue, and zika. A team of experts from stakeholders of health were mobilised to control and/or contain the spread of the disease to other parts of Ghana. The team carried out several activities and assessments to stop the spread of yellow fever. Notably is the investigation to determine different types of mosquitoes involved in transmitting the disease. We collected some mosquitoes and processed them for vital information that could prevent future outbreaks of the aforementioned diseases. There is no surveillance system in Ghana to pick up early warnings regarding potential viral disease outbreaks. Therefore, there is scanty information on these type of mosquitoes and viruses found in different places. We used standard procedures to assess the risk of these mosquitoes in causing future disease outbreaks. Our findings suggested a high risk of future outbreaks for any of the viral diseases tested. We therefore recommended the implementation of a mosquito surveillance system to prevent future outbreaks.

## Introduction

Globally, *Aedes*-borne viruses (ABVs) are arboviruses that are still a major public health concern [1]. About 17% of infectious diseases worldwide are arthropod-borne and significantly the emerging pathogens [2–5]. It has also been estimated that more than 3.9 billion people in Africa are at risk of arboviral diseases [6–8]. In recent times, certain countries in West Africa have experienced outbreaks of arboviruses such as Dengue Fever (DF) and Yellow Fever (YF). These viruses belong to the flavivirus group. They are enclosed RNA viruses and the commonest global human arboviruses (Figueiredo, 2019). In tropical Africa, the yellow fever virus (YFV) affects primates (people and monkeys) [9]. YFV evolved in Africa’s sylvatic cycles, maintained in nonhuman primates and forest-dwelling *Aedes* mosquitoes, and has a history of continuous transmission among humans via *Aedes aegypti* [10]. YFV can cause asymptomatic infection, non-specific clinical disease, and deadly haemorrhagic fevers in humans. Despite the development and presence of a safe vaccine, there was a recent outbreak in Ghana [11]. Other arboviral diseases that have been reported during outbreaks in Africa include Chikungunya fever (CHIKF) and Zika fever (ZIKF) caused by Chikungunya virus (CHIKV) and Zika virus (ZIKV) respectively.

Some authors have reported DF incidence and epidemics in several African countries including Angola (1986, 1999-2002, 2013, 2011-2018), Benin (1987-1993), Cape Verde (2009), Cameroon (1987-1993, 1999-2003; 2006), Comoros (1943-1948, 1984, 1992, 1992-1993), Democratic Republic of the Congo (DRC) (1999-2001, 2007), Djibouti (1991-1992), Egypt (1779, 1887, 1927), Eritrea (2005), Ethiopia (1999-2002, 2007, 2013), Equitorial Guinea 1999-2002), Gabon (1999-2002, 2007), Kenya (1982, 1984-1986), La Réunion (1977, 1978), Madagascar (1943-1948, 2006), Mali (2008), Mauritania (2014-2020), Mauritius (2009), Mozambique (1964-1985, 2014-2015), Namibia (1999-2002, 2006), Nigeria (1964-1968), Rwanda (1987-1993), Senegal (1928, 1979, 1980-1985, 1990, 1999, 2007, 2009-2018), Seychelles (1976-1979), Somalia (1982, 1985-1987, 1992-1993, 2011-2018), South Africa (1927), Sudan (1984-1986), and Tanzania (1987-1993, 1999-2002, 2006, 2010, 2014), Uganda (1999-2002) and Zambia (1987-1993), Zanzibar (1823, 1870, 2010) [12, 13]. Others have indicated that incidence and outbreak of Chikungunya fever (CHIKF) has been reported in several African countries. The first CHIKF outbreak occurred in Tanzania (1952-1953) and Chikungunya virus (CHIKV) was successfully isolated during this period. Between the 1960s and 1980s CHIKV spread and caused outbreaks in countries such as Angola (1970-1971), Brazzaville – Republic of the Congo (2011, 2018), Cameroon (2006), Central African Republic (CAR) (1978-979), Comores (2005-2006), DRC (1958, 1960, 1999-2000, 2003-2012, 2018), Ethiopia (2019), Equitorial Guinea (2002, 2006), Gabon (2007-2010), Kenya (2004, 2014-2016, 2017-2018), La Réunion (2005-2006), Mauritius (2005-2006), Madagascar (2006), Nigeria (1964,1969,1974), Senegal (1960,1997-1998), Seychelles (2005-2006), Sierra Leone (1978), Somalia (2016), South Africa (1956, 1975-1977), Sudan (2005, 2018), Tanzania (2007-2008), Uganda (1961-1962, 1968), Zambia (1959), and Zimbabwe (1957, 1961-1962, 1971) [14, 15]. Additionally, ZIKF had been suggested to be a threat for the whole of Africa. The virus was first found in Uganda (1947) [16]. Since then, its incidence and outbreak have been reported in Angola (2015), Cape Verde (2015), Gabon (2007), Gambia (2007, 2011-2012), Guinea-Bissau (2015), Mali (2007, 2011-2012), Nigeria (1954, 1971), Senegal (2007, 2011-2012), Sierra Leone (1972) [16, 17]. Also, YF incidence and outbreaks reported in Africa include Benin (1996), Ethiopia (1960-1962), Gabon (1994), Gambia (1934-1935, 1978-1979), Ghana (1926-1927, 1937, 1996), Kenya (1992-1995), Liberia (1996), Nigeria (1925-1928, 1986-1991, 1996), Senegal (1748, 1778, 1995-1996), and Sudan (1940) [18].

Ghana is surrounded by countries known to be endemic for dengue [19] and yellow fever. The recent yellow fever outbreak in Togo as well as dengue fever outbreaks in Burkina Faso and Cote d’Ivoire occurred in 2020, 2016 and 2016/2017 respectively [19–21]. Historically, there had also been reports of DF incidence and epidemics in neighbouring countries closest to Ghana i.e. Côte d’Ivoire (1982, 1998, 1999-2002, 2008), Burkina Faso (1925, 1983-1986, 2003-2004, 2007, 2013, 2016-2017), and Togo (1087-1993, 1999-2002) [12]. Yet there has been no report of DF outbreak in Ghana. However, in Ghana, there has been DF detection locally and in travellers (1932, 1987-1993, 2002-2005) [12, 13]. Recently, there has been reports of YF outbreak in Angola (2016), Burkina Faso (1996, 2008), Cameroon (2009, 2010, 2012-2013), CAR (2008-2009), Chad (2013), Côte d’Ivoire (2008, 2010-2011), DRC (2010, 2012-2014, 2016), Ethiopia (2013), Ghana (2012), Guinea (2008-2010), Kenya (2016), Liberia (2009), Nigeria (2017), Republic of Congo (2009), Senegal (2010-2011), Sierra Leone (2009, 2011), Sudan (2012-2013), and Uganda (2011, 2016) [18, 22]. In the year 2021, several African countries had another round of YF outbreak. These countries included Cameroon, CAR, Chad, Côte d’Ivoire, DRC, Gabon, Nigeria, and Republic of the Congo [23]. Ghana in 2021 also had a yellow fever outbreak which started in the Savannah Region and quickly spread to the Upper West, Bono, and Oti regions all in the northern parts of Ghana. In 2022, YF was again reported in countries such as Cameroon, CAR, DRC, Chad, Kenya, Niger, Nigeria, Republic of the Congo, and Uganda [24].

Arthropod-borne viruses are transmitted to people mainly by *Aedes aegypti* in tropical and subtropical areas around the globe [25]. Mosquitoes of the *Aedes* genus are vectors for several emerging diseases that are spreading all over the world [26]. These mosquito species are anthropophilic and endophilic in most parts of the world [27] and they are also invasive. Changing environmental variables and urbanization has contributed immensely to the spread of these *Aedes* mosquitoes [28]. *Aedes aegypti aegypti* have been described as the domestic subspecies that have a preference for human blood, while *Aedes aegypti formosus* is known to be efficient vectors also found in sub-Saharan areas [29] and in forested areas. These characteristics make them significant vectors of arboviruses like dengue fever virus (DENV), yellow fever virus (YFV), zika virus (ZIKV), and chikungunya virus (CHIKV). These vectors could be used as predictors for future outbreaks and hence the need to carry out surveillance. Ghana’s response to the 2021 outbreak was remarkable in the sense that stakeholders of health / experts from diverse fields of study were put together as quickly as possible to handle the menace of the YF outbreak. This research was to carry out the vector study to have information on the risk of infections. Measures were put in place to control the spread and mortality due to YF by the national emergency response team.

Most arboviral infections do not have vaccines and/or medications and are at best managed. Outbreaks of arboviral diseases are therefore a major public health concern globally. It is therefore imperative that we assess the risk of arboviral transmission through an established system of continuous surveillance while waiting for the development of vaccines and/or medications for arboviral diseases. In this commission, we therefore investigated the outbreak of YF in the Savannah Region by assessing the entomological risk after the YF outbreak. The findings of this study give valuable data or the needed information to support public health systems to contain other outbreaks that may occur subsequently within the borders of Ghana and other endemic countries that might experience similar outbreaks in the future.

## Methods

**Figure 1:**
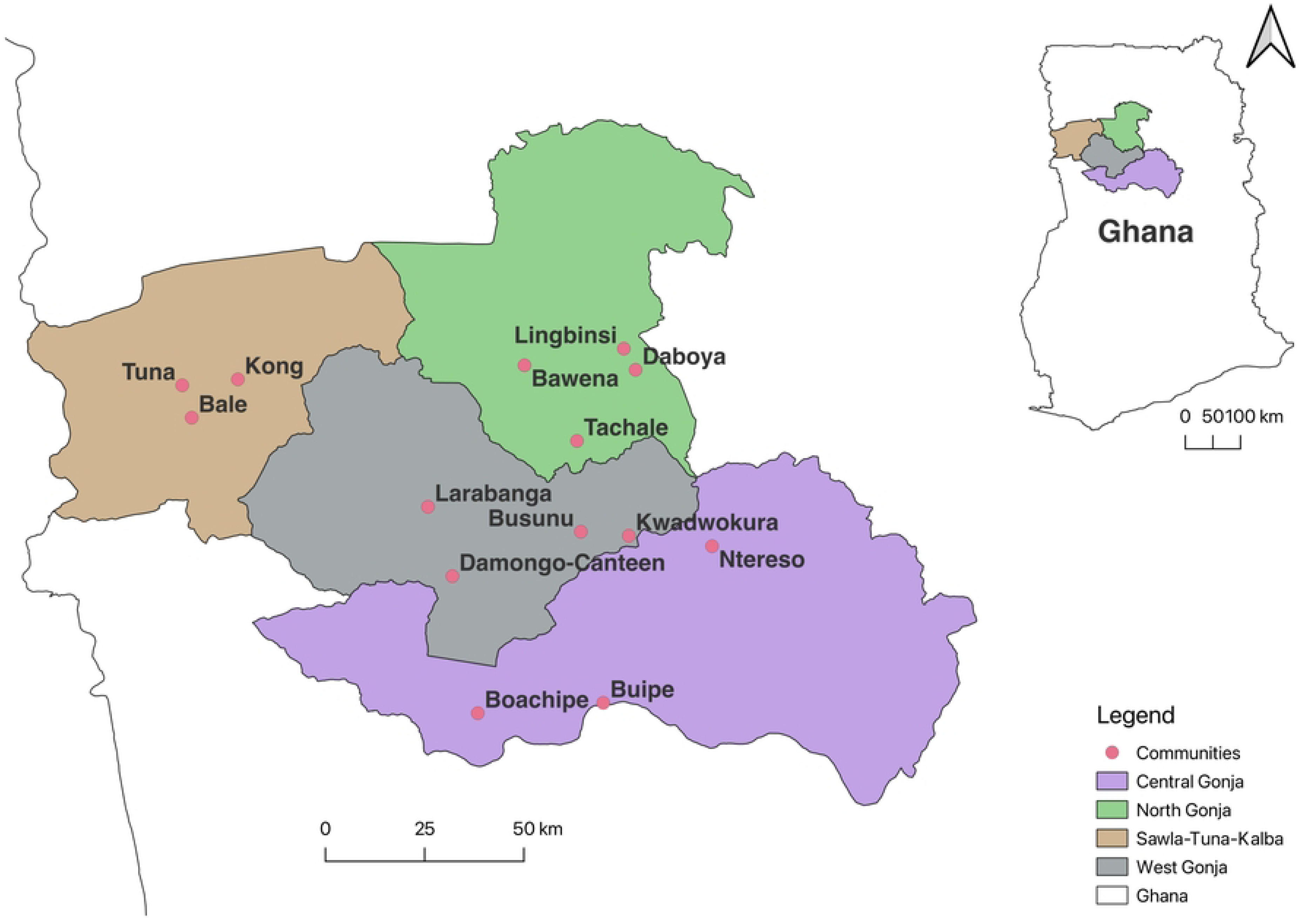
Map of study districts and communities in the Savannah Region of Ghana

The study was conducted in 4 districts of the Savannah Region. These districts included West Gonja, Central Gonja, North Gonja, and Sawla-Tuna-Kalba districts. The Savannah Region lies in the Guinea Savannah vegetation zone and it covers a total area of approximately 35,853 km^2^ making it the largest region in Ghana [30]. This region has a mean annual rainfall and temperature of about 1100 mm and 28.3 °C respectively [31]. The vegetation is mostly grasslands, shrubs and sparsely distributed clusters of drought-resistant trees [30, 31]. The population size was projected to be about 625,299 with a population density of about 17 per km^2^ in the year 2020 [31]. The main occupation in the region is farming.

### Study Design

The Regional Health Directorate (RHD) and the District Health Directorates (DHD) of the Savannah Region were contacted for updates on the outbreak and obtained a list of patients that have been affected in the district (line list). The study districts and communities were then selected based on the information obtained from the line lists. This was followed by community entry / engagements to explain the study to the chiefs, elders, opinion leaders, and members of the selected communities to seek permission for surveillance. This study was a cross sectional survey involving emergency response team moving from one outbreak community to the other. In the sampling of mosquitoes, each of the selected communities in the four districts was divided into 4 imaginary quadrants. Adult and immature stages of *Aedes* mosquitoes were then sampled from each quadrant. Depending on the size of the community, up to 26 houses per quadrant were inspected and sampled for adult and immature stages of *Aedes* mosquitoes.

### Selection criteria

#### Inclusion criteria

The selection of different study districts and communities in the Savannah Region was based on information extracted from the line lists provided by the various DHD. A district was selected based on the number of recorded positive cases from the communities within that district. Between three to four communities per selected district (having reported YF cases) were chosen from the district line list after ranking them based on the reported number of YF positives and death where applicable.

#### Exclusion criteria

Districts and communities having comparatively no or low numbers of YF positives or no death cases associated with YF were excluded from the study.

### *Aedes* mosquito survey

#### Immature *Aedes* mosquito sampling

Water storage containers (categorised into metallic, plastic, clay jar, tire, flowerpot, well, and other water holding objects) were inspected for the larvae and pupae of *Aedes* mosquitoes. Where applicable, about 100 houses in each community were visited to inspect water storage containers for the presence of *Aedes* mosquito larvae and pupae. The number of containers inspected were recorded as either negative or positive for any of the immature stages. The number of houses inspected were also recorded. Other water storage containers in and around houses in the communities were also sampled. All immature stages (larvae and pupae) collected from different quadrant in each community were pooled. These samples were then transported to a makeshift laboratory. It is worth noting that other larval species aside *Aedes* mosquitoes were sorted out. Some of the immature stages were stored in RNAlater while others were reared into adults before storing in RNAlater. The samples were then transported to the Noguchi Memorial Institute for Medical Research. The morphological and molecular species identification was done in the Entomology Laboratory while viral characterisation in mosquito samples (CHIKV, DENV, YFV, and ZIKV) was done in the Virology laboratory

#### Adult *Aedes* mosquito sampling

Three different methods were used for adult *Aedes* mosquito sampling to maximize our chances of getting appreciable numbers. These methods included the use of BG Sentinel Traps, Prokopack Aspirators, and Human Landing Catches (HLC) to collect adult mosquitoes in water storage containers that had adult *Aedes* mosquitoes resting in them or hovering above them. Briefly, mosquito sampling was done by first dividing the community into 4 imaginary quadrants and sampling from each of the quadrants. Ten (10) BG Sentinel traps (2-3 per quadrant) were set each morning in each of the selected communities from 06:00 GMT. These traps were then retrieved at 18:00 GMT on the same day of sampling in each community. HLC was done between the hours of 14:00 GMT to 18:00 GMT in each of the selected communities. The HLC involved 3 collectors per community sampling at various points or suitable locations in the community. Collections involved highly trained members from the emergency response team. These trained collectors always ensured that they do not get bitten when catching mosquitoes. All the vector collectors also had yellow fever vaccination. The Prokopack aspirations were used during inspection of water storage containers for *Aedes* mosquito larvae.

### Species identification and viral characterisation

#### Morphological identification of Aedes mosquitoes

The morphological identification of adult mosquitoes were done using keys as described for *Aedes* [32, 33] and *Culex* [34, 35] .

#### Viral RNA extraction

RNA was extracted from adult mosquito samples from the study sites using QIAGEN RNeasy kit according to manufacturer’s instruction [36] with a few modifications. Each whole mosquito sample was put in a single 1.5 ml microtubes. About 560 µl Buffer AVL containing carrier RNA to a final concentration of 1 µl/µg for this mixture was added to the single mosquito in each microtube. About ten (10) of a 2 mm-diameter beads were added to each microtube. Each mosquito was then homogenised in this mixture for 10 seconds using the BioSpec Products Mini-Beadbeater-96 (BioSpec Products – Catalog No. NC0141170). About 56 µl of the homogenate in each microtube was aliquoted and pooled (up to ten (10) mosquito aliquots / homogenates per pool) to obtain a final volume of 560 µl homogenate per pool. RNA was then extracted following manufacturer’s instructions [36] and eluted to a total volume of 50 μl for viral detection.

#### Detection of arboviruses

RNA was extracted from adult *Aedes* mosquitoes using protocol as described [36]. The extracted RNA was subjected to real time reverse transcription polymerase chain reaction (real time RT-PCR) for the detection of Chikungunya, Dengue, Yellow Fever, and Zika. Briefly, the CDC trioplex master mix and respective viral primers were used for the detection of Chikungunya, Dengue, and Zika viruses using AgPath-ID One-Step RT-PCR kit (Thermo Fisher Scientific, NY, USA) [37–39]. The trioplex detection assay had unique or specific primer sets and probes targeting different regions of the respective viral genomes for the identification of the three different arboviruses mentioned above. For this assay, the nSP1 region was amplified for the detection of Chikungunya using the sense primer (CHIKV-F: 5’-[ACA ATC GGT GTT CCA TCT AAA G]-3’), antisense primer (CHIKV-R: 5’-[GCC TGG GCT CAT CGT TAT T]-3’) and probe (CHIKV-P: 5’-[ACA GTG GTT TCG TGT GAG GGC TAC]-3’); the envelope gene was amplified for Zika virus detection using sense primer (ZIKV-F: 5’-[CCG CTG CCC AAC ACA AG]-3’), antisense primer (ZIKV-R: 5’-[CCA CTA ACG TTC TTT TGC AGA CAT]-3’) and probe (ZIKV-P: 5’-[AGC CTA CCT TGA CAA GCA GTC AGA CAC TCA A]-3’); and the untranscribed region (UTR) of the Dengue virus genome was amplified for the detection of the four serotypes of Dengue using the primer sets (DENV-F: 5’-[TAG TCT RCG TGG ACC GAC AAG]-3’; DENV-R1: 5’-[CAG TTG ACA CRC GGT TTC TC]-3’; DENV-R2: 5’-[GGG TTG ATA CGC GGT TTC TC]-3’) and probe (DENV-P: 5’-[CGY CTW TCA ATA TGC TGA AAC GCG]-3’) . On the other hand, a singleplex master mix was used for the detection of Yellow Fever [37–39]. The Yellow Fever virus was amplified using the sense primer, YFS (5’-[AAT AGT TGC TAG GCA ATA AAC AC]-3’), antisense primer, YFAS (5’-[TCC CTG AGC TTT ACG ACC AGA]-3’), and probe, YFP (5’-[ATT CGT TCG TTG AGC GAT TAG CAG T]-3’) [37–39]. To verify the success of the nucleic acid extraction, a primer set (RP-F: 5’-[AGA TTT GGA CCT GCG AGC G]-3’; and RP-R: 5’-[GAG CGG CTG TCT CCA CAA GT]-3’; and human ribonuclease probe (human Rnase P or RP-P: 5’-[TTC TGA CCT GAA GGC TCT GCG CG]-3’) was added to the master mix for the PCR run [37–39].

### Ethics statement

This was in response to outbreak therefore ethical approval was waived.

### Sample size estimation

Based on the WHO criteria for estimation of risk of arboviral transmission using vector indices, mosquito surveys were conducted in at least 100 houses (about 25 houses per quadrant) depending on the size of the community [40].

## Results

### Environmental and house survey

Total number of houses sampled in all communities selected were 1001 (minimum = 28; maximum = 102). The distribution of the houses in the four (4) districts are as follows; 261 (West Gonja); 197 (Central Gonja); 292 (North Gonja); and 251 (Sawla-Tuna-Kalba). The total number of houses positive for immature stages of *Aedes* mosquitoes were 130. These immature stages of *Aedes* mosquitoes collected after house inspections were 53, 38, 12, and 27 for West Gonja, Central Gonja, North Gonja and Sawla-Tuna-Kalba respectively (Table 1)

**Table 1:**
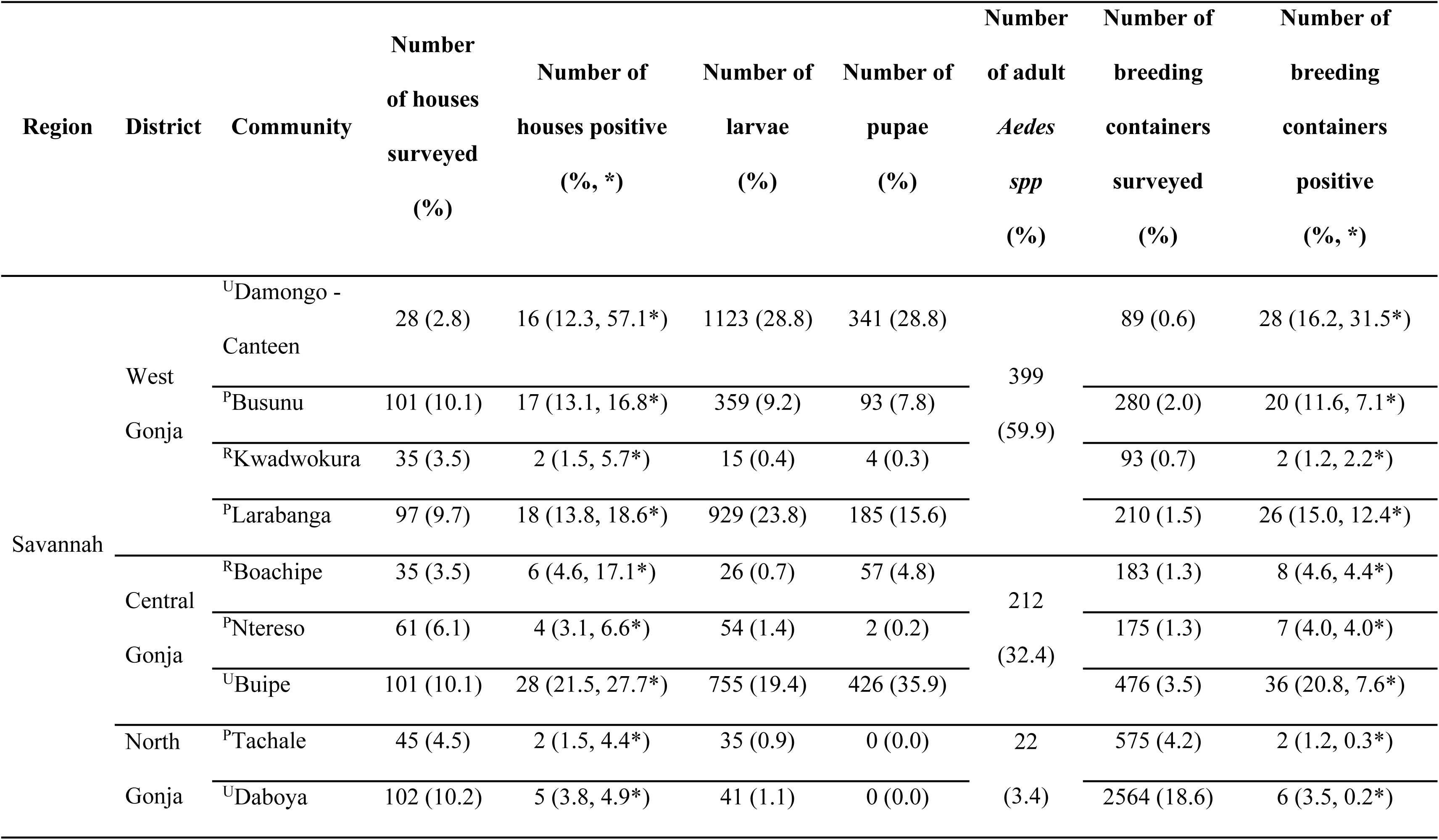
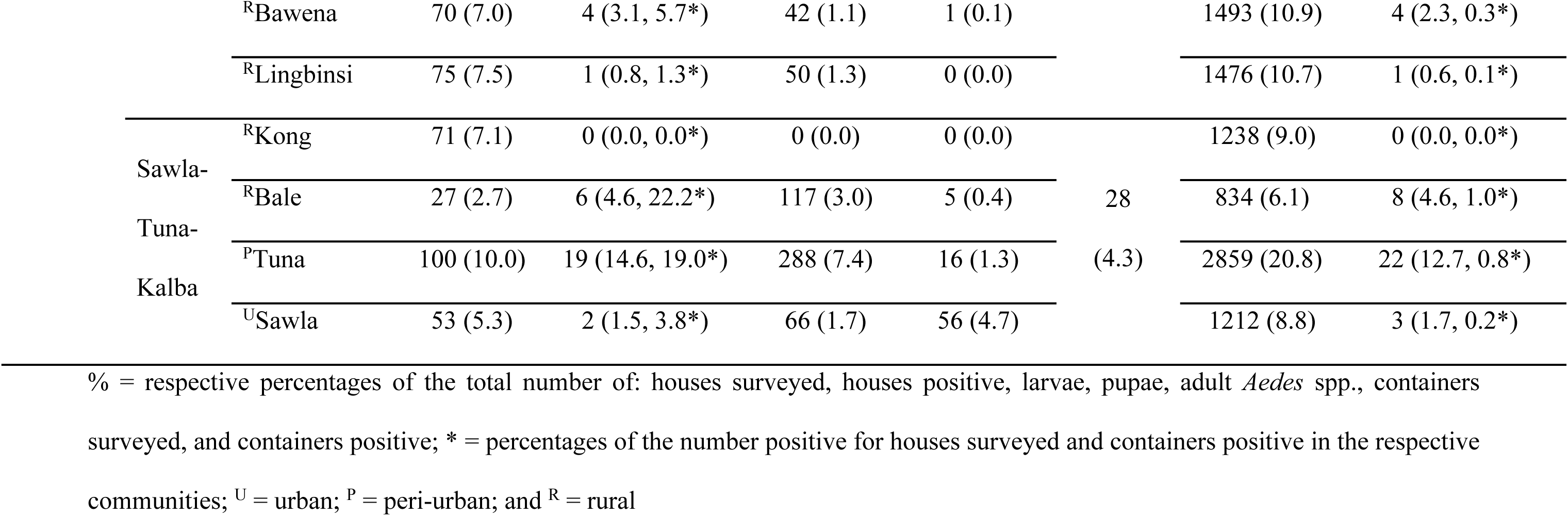
Larval and adult surveys in communities within the districts

**Figure 2:**
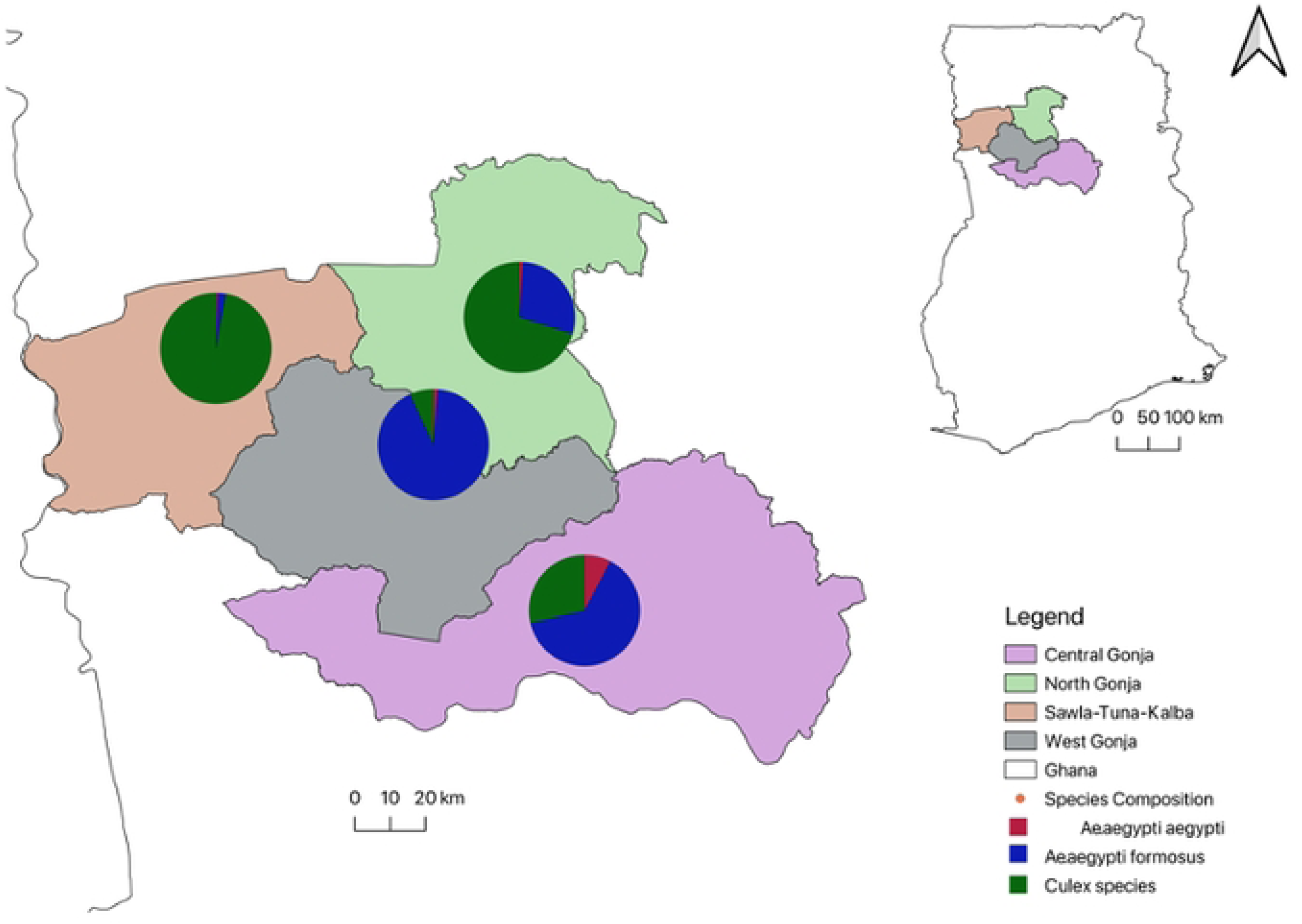
Map showing Mosquitoes caught in each district

### Risk assessments for vector collection

#### Assessment with immature and adult stages of Aedes mosquitoes

While the highest percentages of larvae and pupae were collected from the West Gonja (62.21%, 52.53%) and Central Gonja (21.41%, 40.89%) districts, the highest numbers of breeding containers were found in North Gonja (6108 (44.40%)), and Sawla-Tuna-Kalba (6143 (44.65%)) districts. A greater number of positive breeding containers were, however, found in Central and West Gonja districts (Table 1). In all the study districts, the study revealed that the common breeding containers were metallic and plastic (Table 2)

**Table 2:**
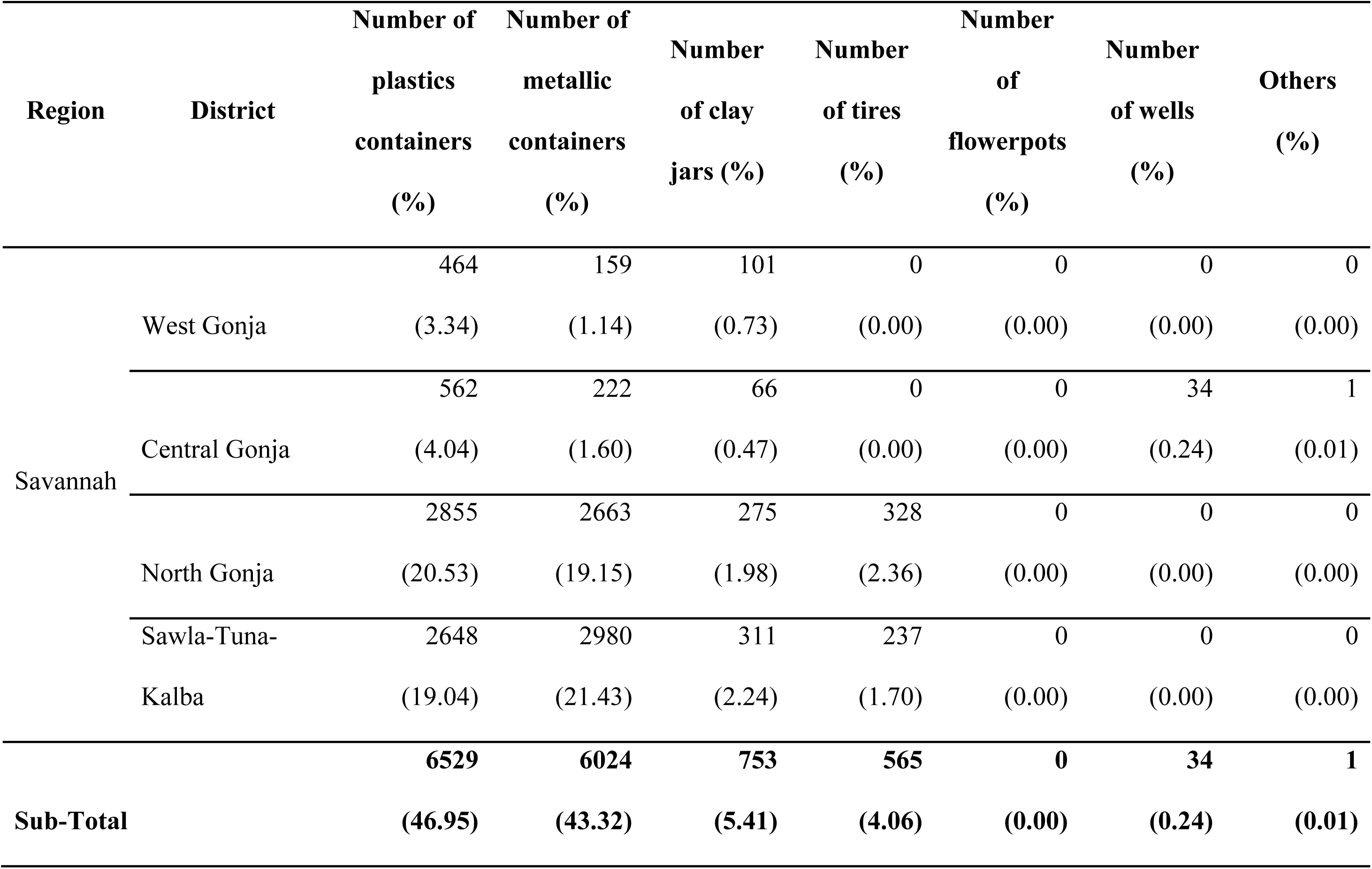
Types of breeding containers surveyed in the study districts

Only two (2) adult *Aedes aegyptii* mosquitoes were caught with the BG Sentinel Trap in West Gonja District. Also, a total of 28 *Aedes aegypti* were collected using Prokopack. No mosquitoes were caught using HLC. All adult *Aedes* mosquitoes (either raised from larval collections or collected with any of the adult mosquito collection methods) were morphologically identified (Table 3).

**Table 3:** Morphological identification of adult mosquitoes from the study districts

| District | Morphological Identification |  | Total (%) |
| --- | --- | --- | --- |
|  | *Aedes aegypti aegypti* | *Aedes aegypti formosus* |  |
|  | Number (%) | Number (%) |  |
| **Central Gonja** | 25 (3.8) | 187 (28.6) | 212 (32.4) |
| **North Gonja** | 4 (0.6) | 18 (2.8) | 22 (3.4) |
| **Sawla-Tuna-Kalba** | 1 (0.2) | 27 (4.1) | 28 (4.3) |
| **West Gonja** | 5 (0.8) | 387 (59.2) | 392 (59.9) |
| **Total** | **35 (5.4)** | **619 (94.6)** | **654 (100.0)** |

In general, there was a high risk of YF and other ABVs transmission in all the communities surveyed in the four districts. However, in Lingbinsi and Kong communities found in North Gonja and Sawla-Tuna-Kalba districts respectively, had entomological indicies that suggested that the risk of YF and other ABVs infections in these communities were low (Table 4).

**Table 4:** Estimation of risk of yellow fever outbreak within communities in the districts

|  | *Aedes* larval indices |  |  | Risk level | Common Types of breeding containers |
| --- | --- | --- | --- | --- | --- |
|  | Container Index (CI) | House Index (HI) | Breteaues Index (BI) |  |  |
| **WHO threshold** | **>3%** | **>4%** | **>5%** |  |  |
| **<u>West Gonja District</u>** |  |  |  |  |  |
| Damongo - Canteen | 23.93 | 57.14 | 3.57 | High risk | Plastic and metallic storage |
| Busunu | 6.67 | 16.83 | 0.20 | High risk | Plastic and metallic storage |
| Kwadwokura | 2.11 | 5.71 | 0.16 | High risk | Plastic and metallic storage |
| Larabanga | 11.02 | 18.56 | 0.28 | High risk | Plastic and metallic storage |
| **<u>Central Gonja District</u>** |  |  |  |  |  |
| Boachipe | 4.19 | 17.14 | 0.65 | High risk | Plastic and metallic storage |
| Ntereso | 3.85 | 6.56 | 0.19 | High risk | Plastic and metallic storage |
| Buipe | 7.03 | 27.72 | 0.36 | High risk | Plastic and metallic storage |
| **<u>North Gonja District</u>** |  |  |  |  |  |
| Tachale | 0.35 | 4.44 | 0.10 | High risk | Plastic and metallic storage |
| Daboya | 0.23 | 4.90 | 0.06 | High risk | Plastic and metallic storage |
| Bawena | 0.27 | 5.71 | 0.08 | High risk | Plastic and metallic storage |
| Lingbinsi | 0.07 | 1.33 | 0.02 | Low risk | Plastic and metallic storage |
| **<u>Sawla-Tuna-Kalba District</u>** |  |  |  |  |  |
| Kong | 0.00 | 0.00 | 0.00 | Low risk | Plastic and metallic storage |
| Bale | 0.95 | 22.22 | 1.10 | High risk | Earthenware pots and metallic storage |
| Tuna | 0.76 | 19.00 | 0.22 | High risk | Plastic and metallic storage |
| Sawla | 0.25 | 3.77 | 0.11 | Low risk | Plastic and metallic storage |

#### Arbovirus detection PCR

Total number of 673 adult mosquitoes consisting of 70 pools were screened for ABVs. Each pool had up to a maximum of 10 *Aedes aegypti* mosquito mixture making up the homogenate used for RNA extractions as describe earlier. The detection of the four arboviruses (CHIKV, DENV, YF, and ZIKV) investigated from pools of *Aedes* mosquito samples were all negative after running real time RT-PCR.

### Data analyses

The WHO has a threshold for estimating risk of arboviral disease transmission using the vector indices. The potential risk for yellow fever outbreak in the study area was assessed using the WHO risk assessment criteria [40]. The respective threshold for Container Index (CI), House Index (HI), and Breteaux Index (BI) are 3.0%, 4.0%, and 5.0%. Beyond these thresholds, the risk of a given arboviral disease outbreak is deemed high. Vector (larval and pupal) indices were estimated using Breteaux Index (BI), Container Index (CI), and House Index (HI). Breteaux index (BI) is defined as the number of containers found positive for larvae and pupae per 100 houses surveyed. The container index (CI) is the percentage of containers that were found positive. The house index (HI) is the percentage of houses found positive for larvae or pupae. In general, there were more *Aedes aegypti formosus* caught during the outbreak than *Aedes aegypti aegypti* (F-test, *p* < 0.05). Using Chi-square test comparison of *Aedes* mosquito species caught during the YF outbreak, it was realised that there was no significant difference between the numbers of *Ae. ae. aegypti* and *Ae. ae. formosus* in Central Gonja and North Gonja (*p* = 0.39), Central Gonja and Sawla-Tuna-Kalba (*p* = 0.19), North Gonja and Sawla-Tuna-Kalba (*p* = 0.09), and Sawla-Tuna-Kalba and West Gonja (*p* = 0.32). However, there was significant difference in the numbers of *Ae. ae. agypti* and *Ae. ae. formosus* caught in Central Gonja and West Gonja (*p* <<< 0.01), as well as North Gonja and West Gonja (*p* <<< 0.01).

### Key findings

A few vital observations were made during the study. Firstly, access to water (pipe-borne, and well) was not much of a problem in the peri-urban (Larabanga) and urban (Buipe, Damango, Daboya, and Sawla) settings, however, these settings had the highest numbers of water storage containers due to unreliable pipe-borne water supply. Secondly, these peri-urban and urban dwellings had highest numbers of water storage containers infested with the immature stages of the *Aedes* mosquitoes. Thirdly, in the rural communities such as Lingbinsi and Kong, water is hardly stored for long periods because water was fetched from streams, rivers, and other nearby water bodies for daily house chores. Lastly, all vectors screened for the four ABVs were negative.

## Discussion

Following a YF outbreak in October 2021 in the Savannah Region of Ghana, an Emergency Response Team (ERT) comprising various sectors of the health system in Ghana and other stakeholders of health was assembled. This ERT was made up of experts from various institutions (including Noguchi Memorial Institute for Medical Research (NMIMR), different department under the Ghana Health Service (GHS), WASH & IPC (Water, Sanitation and Hygiene & Infection Prevention and Control), World Health Organisation (WHO) – Ghana, and Centre for Disease Prevention and Control (CDC) – Ghana) forming several groups such as Laboratory, Environmental, Vector Research, Public Health Action / Response Intervention, Case Management, WASH & IPC, Risk Communication, Community Engagement & Social Mobilisation, Logistics, and Vaccination groups among others. The ERT was tasked to assess, manage, contain and / or control the spread of the YF outbreak. The first case of the YF outbreak was reported on the 15^th^ of October 2021 in the Savannah Region, and it spread quickly to three other regions (Upper West, Bono, and Oti) by 27^th^ of November 2021. Within this period, as many as 202 suspected cases were reported. Out of this number, 70 were confirmed to be YF positives and 35 deaths were reported out of the confirmed cases. The reported Case Fatality Ratio for these study districts was 17%. This study is therefore essential because of the following reasons among others. The Savannah Region of Ghana shares porous borders with neighbouring Cote d’Ivoire and Burkina Faso. The history of arboviral disease outbreaks given in these neighbouring countries suggests that Ghana may share suitable vectors for the transmission of arboviruses with both countries. Therefore, there is the potential for the spread of YF outside of Ghana to these countries following an outbreak. Similarly, in the event of any arboviral outbreak in these neighbouring countries, there is a high potential for the spread of these arboviruses in Ghana. Despite this threat of ABV infections that may cause outbreaks in Ghana periodically, there is no established surveillance systems for ABVs in the country. This study aside solely assessing and proposing measures for containing or controlling the spread of YFV and other possible arboviruses such as CHIKV, DENV, and ZIKV using the vector indices, will provide additional data which will be useful for establishing surveillance systems and planning future interventions in Ghana. Another important component of this study / surveillance is the vector component. Infections leading to outbreaks of YF and other ABV diseases occur when vectors are abundant and efficient in transmitting these ABVs. It is therefore very important that surveillance is conducted during the period of an outbreak. The level of infections is usually high during epidemics. It therefore increases the chances of successfully isolating the ABVs for further studies to determine the lineages and strains of pathogens being transmitted. These are useful information for several vector /entomological, genetic / molecular and immunological / vaccine studies required to treat and / or contain any future ABV outbreaks. Conducting post outbreak surveillance periodically would also help avert potential ABV outbreaks or efficiently control these outbreaks whenever they occur in the future. The *Aedes* mosquito populations found in endemic areas, their vector biology, and respective densities among others would be known for the right measures and / or interventions to be put in place to effectively and efficiently handle any future outbreaks.

*Aedes* mosquitoes are known to inhabit closely to humans [41]. They usually breed in water-holding containers including clay pots, metallic and plastic containers, discarded car tires, and other containers both inside and outside human houses [42]. Environmental and house surveys were therefore conducted for immature (larvae and pupae) and adult *Aedes* mosquitoes. While a few containers were found within the community serving as breeding sites, many water storage containers (mostly different types of metallic, plastic, and clay jar containers) and / or *Aedes* mosquito breeding sites were found in the different houses visited (Tables 1 & 2). This therefore increases the domestic *Aedes* breeding sites and hence the man-vector contact rates due to proximity of these arboviral vectors that infect humans. The major risk factors associated with the emergence of Yellow Fever (YF) and other arboviral diseases (including Chikungunya, Dengue, and Zika) were domestic water storage containers that were suitable for vector development (Table 1). This observation agrees with findings from work done by Ouattara and colleagues [42]. In all four (4) districts, communities (urban, peri-urban, and rural) having reliable water supply and / or potable water sources tend to have lower numbers of water storage containers (Table 1). Also, it was observed that communities / districts having wells as one of their sources of water had high number of water storage containers (Tables 1 & 2). It was worth noting that there were more positive breeding containers in the urban and peri-urban settings than the numbers recorded in the rural settings. This is because most of these rural communities have streams or rivers from which they get water for their daily activities. Most households in the rural settings therefore may not have the need to store water for longer periods to serve as breeding sites for these *Aedes* mosquitoes. There was therefore a correspondingly lower numbers of immature stages (larvae and pupae) of *Aedes* mosquitoes obtained from breeding containers in the rural areas compared to the urban and peri-urban settlements (Table 1). When assessed with the WHO thresholds for entomological indices, this translated into high risk of ABV transmission should there be an outbreak in these study areas (Table 3).

In all the communities (urban, peri-urban, and rural), the only Aedine populations obtained from the larval surveys conducted were *Aedes aegypti aegypti* and *Aedes aegypti formosus*. The *Ae*. *ae*. *aegypti* population has adapted well to urban settings and responsible for most ABV transmissions in these areas while *Ae*. *ae*. *formosus* are noted for ABV transmission in forested areas and therefore important in maintaining sylvatic and intermediate YF transmissions in primates and humans respectively. The predominant ABV vector found during the YF outbreak in all study districts was *Ae*. *ae*. *formosus* suggesting that they might be the *Aedes* population driving the transmission and quick spread of YF during this epidemic. In this outbreak, YF cases were reported mostly from nomadic populations who had moved from Nigeria into a forest reserve in Ghana’s Savannah region which is visited by tourists. Reports from the GHS indicated vaccination coverage within the nomads being very low. In Ghana YF is endemic, and it is associated with severe disease in approximately 15% of cases. Notwithstanding the fact that there is high overall population immunity against YF in Ghana (88% in 2020 according to WHO-UNICEF estimates), there are pockets of the Ghanaian population that are not vaccinated, including unvaccinated nomadic people. Most of these individuals during this survey lived usually close to the forest, middle of the forest (particularly the nomadic populations), and other areas difficult to reach during vaccination campaigns. These two populations remain at risk for YF (and other arboviral diseases) and could be responsible for the maintenance / continued YF transmission. A routine YF vaccination campaign is done to vaccinate the unvaccinated populations. The current outbreak investigation found settlements of newcomer populations who had arrived after the last mass campaign and were largely unvaccinated (Source:https://www.who.int/emergencies/disease-outbreak-news/item/yellow-fever---ghana). There is therefore the potential for a rapid spread of YF within such unvaccinated populations considering that most of these vulnerable populations live in forested areas or close to the edge of forests making it easy for them to suffer sylvatic arboviral spillover infections [43]. Having high densities of efficient vectors such as the *Aedes aegypti* populations in the study districts, arboviral infections in the sylvatic cycle could re-emerge leading to an outbreak [27, 43].

This study revealed a very high number of *Aedes aegypti formosus* (Table 4). This is the vector known to maintain the savannah YF transmission cycle after picking the infection from the primates of the sylvatic (forest) transmission cycle. In close proximity to unvaccinated people, these vectors when infective would transmit the YF virus to these individuals upon taking blood meal leading to a possible epidemic or outbreak. It was therefore imperative, during the arboviral risk assessment for the YF outbreak, to ascertain the risk of arboviral infections in the study districts particularly within settlements of these vulnerable human populations using the entomological indices.

Entomological indices are recommended by the World Health Organization for monitoring vector density and evaluating control measures [44]. The house index HI, which measures the percentage of houses infested with larvae or pupae; the container index (CI), which measures the percentage of water-holding containers infested with larvae or pupae; and the Breteau index (BI), which measures the number of positive containers per 100 houses inspected, are all related to immature mosquito populations [45]. In general, our arboviral risk estimation revealed a high risk for the transmission of YF and other arboviruses in all the study districts (Table 3). This may be affirmed by the presence of the arboviral vectors in the study districts, suitable vector breeding containers found in and around houses, habitual water storage practices usually in the urban communities due to unreliable supply of potable water, and water storage containers mostly uncovered and when / where covered may usually have broken lid(s) allowing access for these mosquito vectors to use them as breeding containers. The appreciably high densities of the arboviral vector populations found in the study areas also support this risk evaluation especially in the West and Central Gonja districts where relatively large numbers of the *Aedes* vector populations were found. One of the major occupations in these study districts is farming. The vegetation is predominantly savannah. Most of these farmlands are found in and around savannah and forested areas where most of the *Aedes aegypti formosus* responsible for maintaining YF transmission in primates (monkeys and humans) usually reside in abundance breeding in rock pools and tree axils among others. Farming and other human activities such as hunting, hiking, camping, and construction of new houses has brought humans close to the edge of forested lands and areas where these vectors thrive. This may therefore increase man-vector contacts for those farmers having their farms in such locations. Though in low numbers, *Aedes aegypti aegypti* known to maintain the urban YF transmission was also found in every district. These two *Aedes aegypti* populations together can concurrently spread arboviral diseases at a very fast pace during an outbreak. None of the four arboviruses namely CHIKV, DENV, YFV, and ZIKV was detected after running real-time RT-PCR. During an outbreak, it is usually expected that with up to a few hundreds of these mosquitoes screened, a successful detection of any of these arboviruses could be realised. Although no arbovirus was detected molecularly with RT-PCR, data from our study suggest the two Aedine populations played vital roles in the spread of the YFV during the outbreak of YF in the Savannah Region of Ghana with the principal vector being *Aedes aegypti formosus*. There may be a few reasons for these zero detections of YF and the other arboviral diseases. Inadequate numbers of *Aedes* mosquitoes were caught with all the adult collection methods. The adult mosquito collection methods used during *Aedes* mosquito sampling included human landing catches (HLC), BG Sentinel Traps, and Prokopack collections. These three adult mosquito sampling methods was therefore used to maximize the number of the adult Aedine population directly caught from the endemic areas experiencing the outbreak for arboviral detection. The adult *Aedes* mosquitoes targeted or sampled during arboviral disease outbreaks is to allow the investigators capture a sizeable proportion of the adult vector population(s) actively infecting individuals and spreading the disease during this period. However, the total adult *Aedes* mosquitoes directly caught during the YF outbreak using the three adult *Aedes* collection methods was woefully low. For instance, no adult mosquito was caught using HLC, and only 30 *Aedes* mosquitoes were caught with the other two methods (BG Sentinel Trap = 2 and Prokopack collections = 28). The other adult mosquitoes screened for YF and the other arboviruses (CHIKV, DENV, and ZIKV) were all raised from the immature stages (larvae and pupae). Secondly, the nomad population settle in forested areas away from the main communities. They, however, frequent the main communities for some supplies vital to their survival. The *Aedes aegypti formosus* population are also known to thrive and maintain YF transmission in the forested areas. The settlements of the nomads located in the forested areas place them in close proximity to these *Ae. ae. formosus* population and hence increasing nomad-vector contact. During the YF outbreak, it was realised that most of the patients were nomads. These nomads are largely unvaccinated because they move around a lot from one settlement to another and therefore hardly get the opportunity to participate in vaccination programmes. They therefore served as the main source of infections for the YF transmission / outbreak. Even if initially, there were no YF infections among the nomad populations infective *Ae. ae. formosus* mosquitoes could quickly spread YF infections to these unvaccinated nomads. In the main communities, there are pockets of individuals who have not been vaccinated against YF so when infected nomads bring the YF infections to the main communities only these individuals are most likely to be infected. The large, vaccinated populations in the communities therefore do not allow *Aedes* mosquitoes pick up YF infections when taking bloodmeal from individuals in the communities hence a higher proportion of these mosquitoes remain unifected. Thirdly, there are insect-specific viruses known to prevent the development of arboviruses in mosquitoes. For instance, Ghana has similar vectors for transmitting arboviruses such as all the serotypes of DENV in neighbouring Burkina Faso, Côte d’Ivoire, and Togo yet DENV outbreak has never occur in Ghana before. Research conducted prior to the 2021 YF outbreak revealed insect-specific viruses in these vectors after molecular analysis [46]. This could possibly explain to a large extent why it is difficult detecting arboviruses in *Aedes* mosquitoes caught and screened periodically.

## Conclusion

Yellow fever is endemic in Ghana even though about 88% of Ghanaian have been vaccinated against this virus. YFV was not detected in the mosquito samples collected from the study districts, however, *Aedes aegypti* populations were the vectors that spread the YFV during the YF outbreak. Based on the data collected during the study, *Aedes aegypti formosus* is the principal vector for the transmission of arboviruses in the study districts. And the risk of arboviral disease transmission in the study districts is woefully high.

## Recommendation

It will be an incentive for members of any of the study community to empty all abandoned containers filled with water to help reduce the breeding sources for the arboviral vectors. All storage containers filled with water must be tightly covered and emptied if these containers have no covers and / or if the water stored will not be used in a relatively short time. This is also to reduce the number of breeding sources for arboviral vectors. However, to prevent or reduce the chances of arboviral transmission in the study districts or any endemic area due to the estimated high risk of arboviral transmission, it is advisable for the indigenes to get vaccinated. Residents in these high-risk districts are also advised to wear long clothing to cover up exposed skin to prevent or minimise man-vector contacts.

## Acknowledgements

We are very grateful for the support given by the Regional Health Directorate led by the Regional Director of Health, Dr Chrysantus Kubio and all District Health Directorate Teams in North, West and Central Gonja and Sawla-Tuna-Kalba. We also appreciate the support of the Savannah Regional Health Research Officer, Mr. Ibrahim Abubakari, the Savannah Regional Disease Control Officers and all the District Disease Control Officers of Central Gonja, North Gonja, West Gonja, and Sawla-Tuna-Kalba districts. We thank the people, opinion leaders, elders, and chiefs of the study districts for allowing us into their households and giving us permission to conduct the study in their communities. We appreciate the support from personnel in the various Community Health Planning and Services (CHPS) compounds we engaged during this study. We also thank all the staff of the Noguchi Memorial Institute for Medical Research who contributed to this study in diverse ways.

## Authors’ contributions

Conceptualization: SKD; data curation: JHNO, MO, HAB, JAO, SCAA, ROTM, CNLMT, SOA, SC, and MCE; formal analysis: SOA, AAA, SPB, KKF, MO, HAB, and JHNO; funding acquisition: SKD, investigation: DP, RB, BA, SOA, AAA, MA, MCAAK, SPB, KKF, and JHNO; methodology: SKD, MAA^†^, DAB, JHKB, SNP and CK; project administration: SKD; resources: SKD, MAA^†^, DAB, JHKB, and SNP; supervision: SKD, MAA^†^, DAB, JHKB, and SNP; validation: DP, RB, BA, SOA, AAA, MA, MCAAK, SPB, KKF, and JHNO; visualization: SKD, MAA^†^, DAB, JHKB, SNP and JHNO; writing – original draft: SKD, MAA^†^, DAB, SNP, JHKB, and JHNO; writing – review & editing: all authors.

## Funding

We also acknowledge the financial support given by NAMRU-3 under the Global Emerging Infections Surveillance and Response System (GEISS), Japan Agency for Medical Research and Development (AMED), Japan and the Center for Disease Control and Prevention (CDC, Atlanta) through the West African *Aedes* Surveillance Network (WAASuN).

## Conflict of interest

The authors of this study declare no conflict of interest.

## Notes

### Competing Interest Statement

The authors have declared no competing interest.

